# Developmental PCB exposure disrupts synaptic transmission and connectivity in the rat auditory cortex, independent of its effects on peripheral hearing threshold

**DOI:** 10.1101/2020.07.28.224048

**Authors:** Christopher M. Lee, Renee N. Sadowski, Susan L. Schantz, Daniel A. Llano

**Author notes:** Corresponding Author: Daniel A. Llano, 2355 Beckman Institute, 405 N Mathews Ave, Urbana, IL 61801.

## Abstract

Polychlorinated biphenyls (PCBs) are enduring environmental toxicants and exposure is associated with neurodevelopmental deficits. The auditory system appears particularly sensitive, as previous work has shown that developmental PCB exposure causes both hearing loss and gross disruptions in the organization of the rat auditory cortex. However, the mechanisms underlying PCB-induced changes are not known, nor is it known if the central effects of PCBs are a consequence of peripheral hearing loss. Here, we study changes in both peripheral and central auditory function in rats with developmental PCB exposure using a combination of optical and electrophysiological approaches. Female rats were exposed to an environmental PCB mixture in utero and until weaning. At adulthood, auditory brainstem responses were measured, and synaptic currents were recorded in slices from auditory cortex layer 2/3 neurons. Spontaneous and miniature inhibitory postsynaptic currents (IPSCs) were more frequent in PCB-exposed rats compared to controls and the normal relationship between IPSC parameters and peripheral hearing was eliminated in PCB-exposed rats. No changes in spontaneous EPSCs were found. Conversely, when synaptic currents were evoked by laser photostimulation of caged-glutamate, PCB exposure did not affect evoked inhibitory transmission, but increased the total excitatory charge, the number and distance of sites that evoke a significant response. Together, these findings indicate that early developmental exposure to PCBs causes long-lasting changes in both inhibitory and excitatory neurotransmission in the auditory cortex that are independent of peripheral hearing changes, suggesting the effects are due to the direct impact of PCBs on the developing auditory cortex.

**Significance Statement:** The mechanisms by which developmental exposure to polychlorinated biphenyls (PCBs) disrupt the central nervous system are not yet known. Here we show that developmental PCB exposure is associated with long-lasting dysregulation of both excitatory and inhibitory neurotransmission in the rodent brain. We further find that, unlike controls, synaptic parameters in the auditory cortex of PCB-exposed rats are independent of peripheral hearing changes. These data suggest that PCB-related changes in the auditory cortex are independent of their effects on the auditory periphery and that PCB exposure may disrupt the plastic mechanisms needed to restore normal processing in the auditory cortex after peripheral hearing loss.

## Introduction

Polychlorinated biphenyls (PCBs) are a family of compounds originally manufactured for many applications, including dielectrics, hydraulic fluids, coolants, and lubricants. PCBs are composed of a biphenyl molecule with chlorine substitutions at any of the ten positions on the biphenyl molecule, creating up to 209 possible congeners. The physical properties and biological effects of PCBs depend on the positions and number of chlorine substitutions. Although their manufacture in the US was banned in 1978, they persist in the environment, and bioaccumulate and biomagnify in food chains, especially in aquatic species, due to their resistance to degradation and their lipophilicity. Additionally, PCBs are transferred to the fetus and infant through the placenta and breast milk (Agency for Toxic Substances and Disease Registry, 2000; for review: Crinnion, 2011).

Humans and rodents exposed to PCBs experience auditory dysfunction, including higher sound detection thresholds (Goldey et al., 1995; Grandjean et al., 2001; Powers et al., 2006; Trnovec et al., 2008; Min et al., 2014; Li et al., 2015), loss of outer hair cells (Crofton et al., 2000), reduced otoacoustic emission amplitudes (Lasky et al., 2002; Powers et al., 2006; Trnovec et al., 2008), and increased susceptibility to and severity of audiogenic seizures (Poon et al., 2015; Bandara et al., 2016). Complex auditory behaviors such as precise sound localization (Lomber and Malhotra, 2007), temporal processing (Threlkeld et al., 2008), and frequency discrimination of complex stimuli (Znamenskiy and Zador, 2013) require auditory cortical processing in mammals. Developmental exposure to PCBs alters the physiology of the auditory cortex, including delayed auditory P300 latencies (Vreugdenhil et al., 2004), disrupted tonotopic organization of receptive fields (Kenet et al., 2007), and increased sensitivity to GABA blockade (Sadowski et al., 2016). However, the synaptic mechanisms underlying these changes are not known. In addition, it is unclear to what degree these changes are due to direct actions of PCBs in the brain, or whether these changes are secondary effects of peripheral hearing loss.

Hearing loss, when experimentally induced by high level sound exposure, cochlear ablation, or administration of an ototoxic agent, drives plasticity in central auditory structures, weakening inhibitory connections, strengthening excitatory connections, and increasing excitability and spontaneous firing, and these changes generally occur over the course of weeks (Bledsoe et al., 1995; Wang et al., 2002; Vale and Sanes, 2002; Kotak et al., 2005; Sarro et al., 2008; Yang et al., 2012; Chambers et al., 2016; Balaram et al. 2019). These changes effectively increase the gain of the central auditory system and may serve a homeostatic role in restoring central auditory processing after a loss of sensory input (Noreña, 2011; Zeng, 2013; Chambers et al., 2016). Because PCB exposure elevates hearing thresholds, the central auditory system might be expected to respond by reducing inhibition and increasing gain in central structures. Consistent with these predictions, PCB exposure reduces expression of GAD65 in the inferior colliculus (Bandara et al., 2016). However, in the cortex, GAD65 levels are unaffected and thalamocortical transmission is more vulnerable to GABA antagonism in PCB-exposed rats, suggesting PCB treatment is associated with paradoxically higher background levels of cortical inhibition (Bandara et al., 2016; Sadowski et al., 2016).

Synapses in the supragranular layers of the auditory cortex connect neural circuits responsible for a wide range of auditory processes, including cross-frequency integration, sensory gain, coincidence detection, and cross-modality integration (Winkowski and Kanold, 2013; Kato et al., 2015; Jiang et al., 2015; Meng et al., 2017). Therefore, it is important to examine whether PCB exposure affects synaptic connectivity and transmission, as changes could point to underlying causes of complex auditory deficits. Layer 2/3 neurons receive thalamic input, and cortical inputs from all layers, but are more likely to be connected to nearby inputs from layers 2-4 (Oviedo et al., 2010; Atencio and Schreiner, 2010).

To determine the effects of developmental PCB exposure on cortical synaptic transmission, and whether these changes are related to peripheral hearing loss, we dosed rats with either a 6 mg/kg/day PCB oil mixture or a control oil mixture beginning four weeks before breeding and continuing until weaning. Because the properties of PCBs vary among congeners, we studied the effects of an environmentally relevant PCB congener mixture. Experimental subjects were treated with a PCB mixture that mimics the congener profile found in the Fox River in Wisconsin (Kostyniak et al., 2005). From adult offspring, we recorded auditory brainstem responses, and excitatory and inhibitory synaptic currents from layer 2/3 auditory cortical neurons, either in the absence of stimulation (spontaneous and miniature currents), or during laser photostimulation of caged glutamate (evoked currents).

## Methods

### PCB exposure and breeding

All procedures were approved by our university Institutional Animal Care and Use Committee. Rats were maintained in facilities accredited by the Association for the Assessment and Accreditation of Laboratory Animal Care. All animal handling and data collection were performed by experimenters blinded to treatment group. Experimental design and dosing and breeding time courses are summarized in Figure 1. Long-Evans rats, 8-10 weeks of age and of both sexes, were purchased from Envigo, and individually housed in standard polycarbonate cages with woodchip bedding. All rats were fed rat chow (Envigo Teklad rodent diet 8604) and water ad libitum. Females were randomly assigned to control or experimental treatments. Beginning one week after receiving the rats, experimental subjects were orally dosed with a PCB mixture in a corn oil vehicle (6 mg/kg/day PCB mixture) and control subjects were orally dosed with corn oil alone (0 mg/kg/day PCB mixture). Dosing was accomplished by pipetting the PCB mixture or oil (0.4 mL/kg) onto one half of a vanilla wafer cookie (Keebler Golden Vanilla Wafers), which were fed to the rats each day. The PCB mixture (35% Aroclor 1242, 35% Aroclor 1248, 15% Aroclor 1254, 15% Aroclor 1260) was synthesized to mimic the congener profile found in the walleye fish from the Fox River in Wisconsin (Kostyniak et al., 2005). Experimental rats were dosed at 6 mg/kg/day, as developmental exposure at this concentration is ototoxic, and increases audiogenic seizure incidence and severity, but does not produce overt signs of clinical toxicity (Kostyniak et al., 2005; Powers et al., 2006, 2009; Bandara et al., 2016). After four weeks of PCB exposure, each female rat was paired with an untreated male rat in a hanging wire cage. Upon detection of a sperm plug indicating gestational day 0, females were removed from males and daily PCB or control dosing continued through gestation and nursing, until weaning. Litters were standardized to 8 pups two days following birth (PND 2), and pups were weaned on PND 21. All offspring were housed in pairs or triplets with cagemates of the same sex and same treatment. All data presented in the current study were collected from female subjects (PCB: n = 60, control: n = 59).

**Figure 1.**
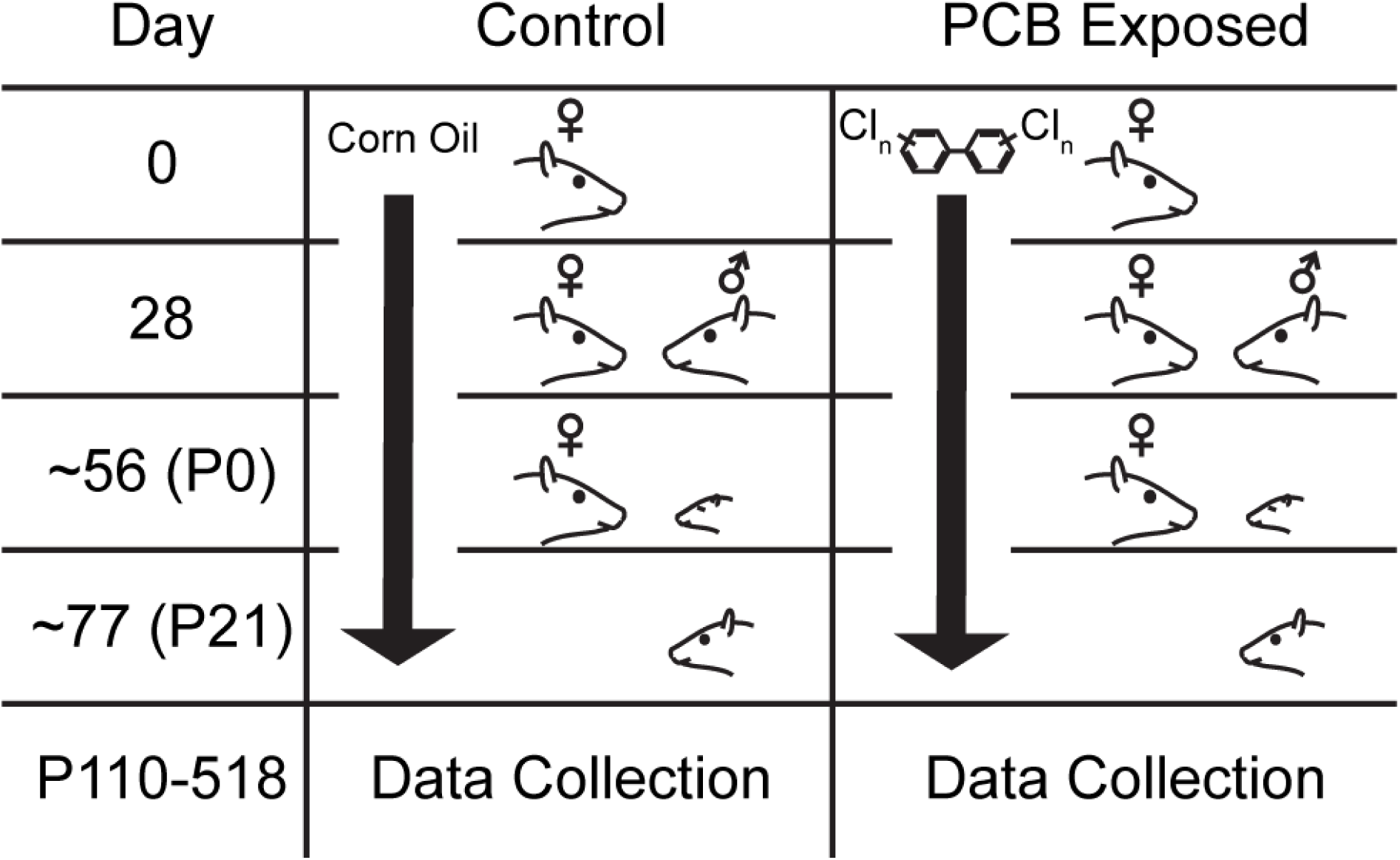
Experimental design and summary timeline of PCB treatment Rows indicate significant experimental timepoints: beginning of dosing (day 0), pairing with male (day 28), parturition (approx. day 56), and weaning (approx. day 77).

Rats were dosed and bred from 5/13/2015 to 8/9/2015 by RNS and were used to collect spontaneous and miniature excitatory and inhibitory postsynaptic potentials. A second group of rats was dosed and bred from 10/10/2016 to 1/4/2017 by CML and was used to collect input maps by laser photostimulation. Auditory brainstem responses (ABRs) were collected from all rats to measure differences in hearing thresholds between treated and control rats.

### Auditory Brainstem Responses

ABRs were collected within one week before electrophysiological recording experiments. Rats were anesthetized with ketamine (100 mg/kg) and xylazine (3 mg/kg) and placed in a sound-attenuated chamber. White noise bursts and pure tone pips, both of 5 ms duration, were delivered through an electrostatic speaker (ES1, Tucker Davis Technologies) placed 2.5 cm from the right ear. ABRs were recorded with two subdermal recording electrodes, one placed above the vertex of the skull, one placed behind the right pinna, and one subdermal ground electrode placed at the base of the tail. The electrodes were connected to a 2400A extracellular preamplifier and headstage (Dagan Corporation), or a RA16PA preamplifier and collected on an RP2.1 real-time processor (Tucker Davis Technologies). Signals were digitized and averaged across 512 trials, and bandpassed between 50 and 3000 Hz. ABR thresholds were estimated as the lowest sound level producing a peak in the signal at 3-5 ms following the sound onset.

### Electrophysiology

We investigated synaptic inputs to auditory cortex of both hemispheres with patch clamp electrophysiology in cortical slices. Rats were anesthetized with ketamine and xylazine, and transcardially perfused with an ice-cold high-sucrose solution (in mM: 206 sucrose, 26 NaHCO_3_, 11 glucose, 10 MgCl_2_, KCl 2.5, NaH_2_PO_4_ 1.25, CaCl_2_ 0.50). The brain was quickly removed and 300 μm thick coronal slices were prepared and allowed to incubate in an oxygenated incubation solution for 1 hour at 32°C. Slices containing auditory cortex were identified based on visual comparison to a mouse brain atlas (Paxinos and Franklin, 2004). After incubation, a slice was transferred to a recording chamber and immersed in an oxygenated artificial cerebrospinal fluid (aCSF, in mM: 126 NaCl, 26 NaHCO_3_, 10 glucose, 2.5 KCl, 2 CaCl_2_, 2 MgCl_2_, 1.25 NaH_2_PO_4_) at 32°C. Pyramidal neurons in layer 2/3 were visualized under DIC microscopy, and whole-cell configuration was achieved with borosilicate glass recording pipettes (pipette resistances of 4-10 MΩ) filled with internal solution (in mM: 117 CsOH, 117 gluconic acid, 11 CsCl, 1.0 MgCl_2_, 0.07 CaCl_2_, 10 HEPES, EGTA 0.1, 2.0 Na-ATP, 0.4 Na-GTP; with pH 7.3). Fig. 3A depicts the location of the recording pipette in an example auditory cortical slice. Electrophysiological signals were sampled at 20 kHz on a DigiData 1550A A/D converter (Molecular Devices).

**Figure 2.**
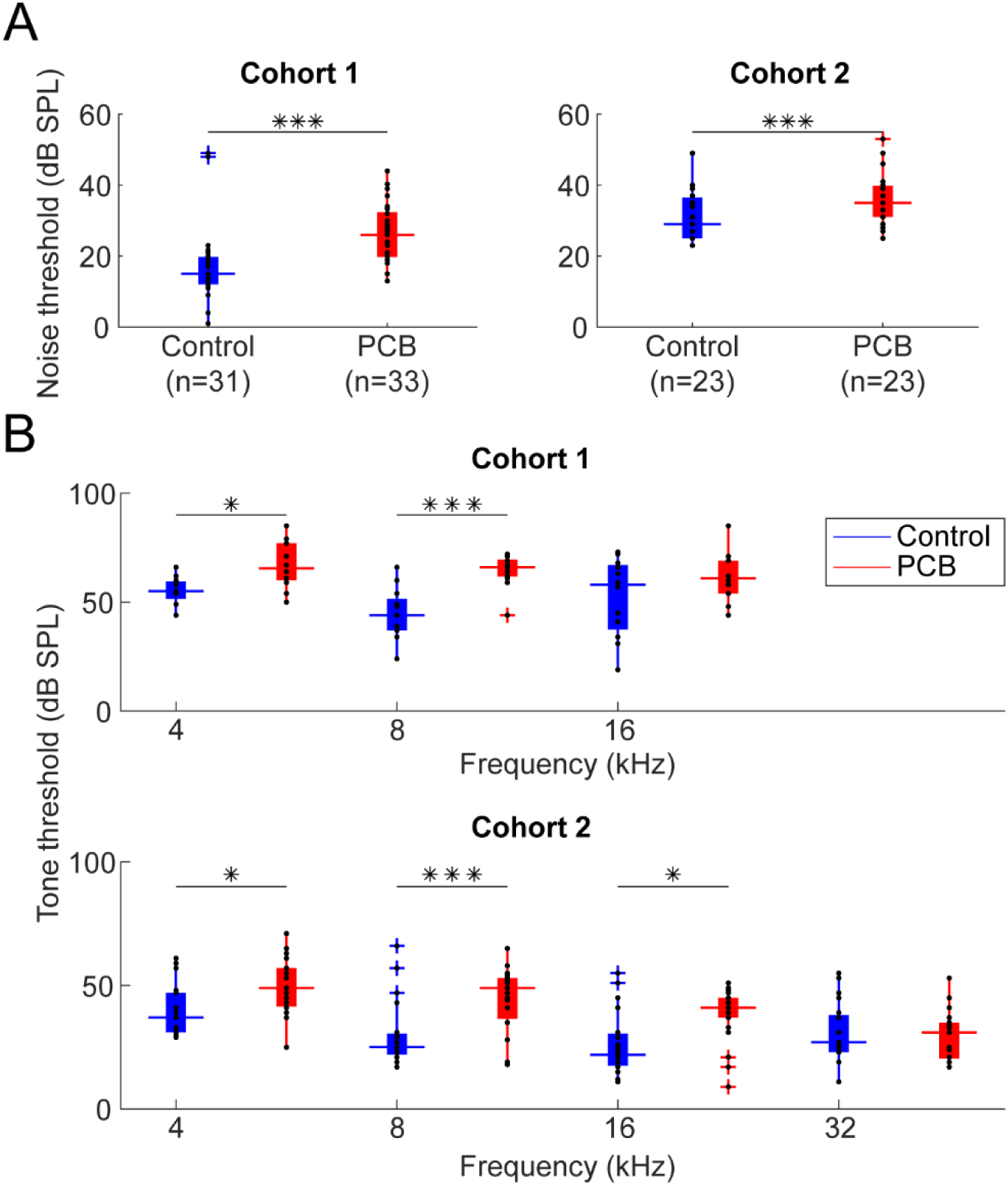
Comparison of ABR thresholds A. Comparison of ABR thresholds in response to noise between control (blue) and PCB (red) treatments. Boxplots indicate median (horizontal bar), 25^th^ and 75^th^ percentiles (box), range of non-outlier points (vertical whiskers), and outliers (crosses). Black asterisks indicate significant comparisons, * p < 0.05, *** p < 0.001. B. Comparison of thresholds to 4, 8, 16, and 32 kHz tones.

**Figure 3.**
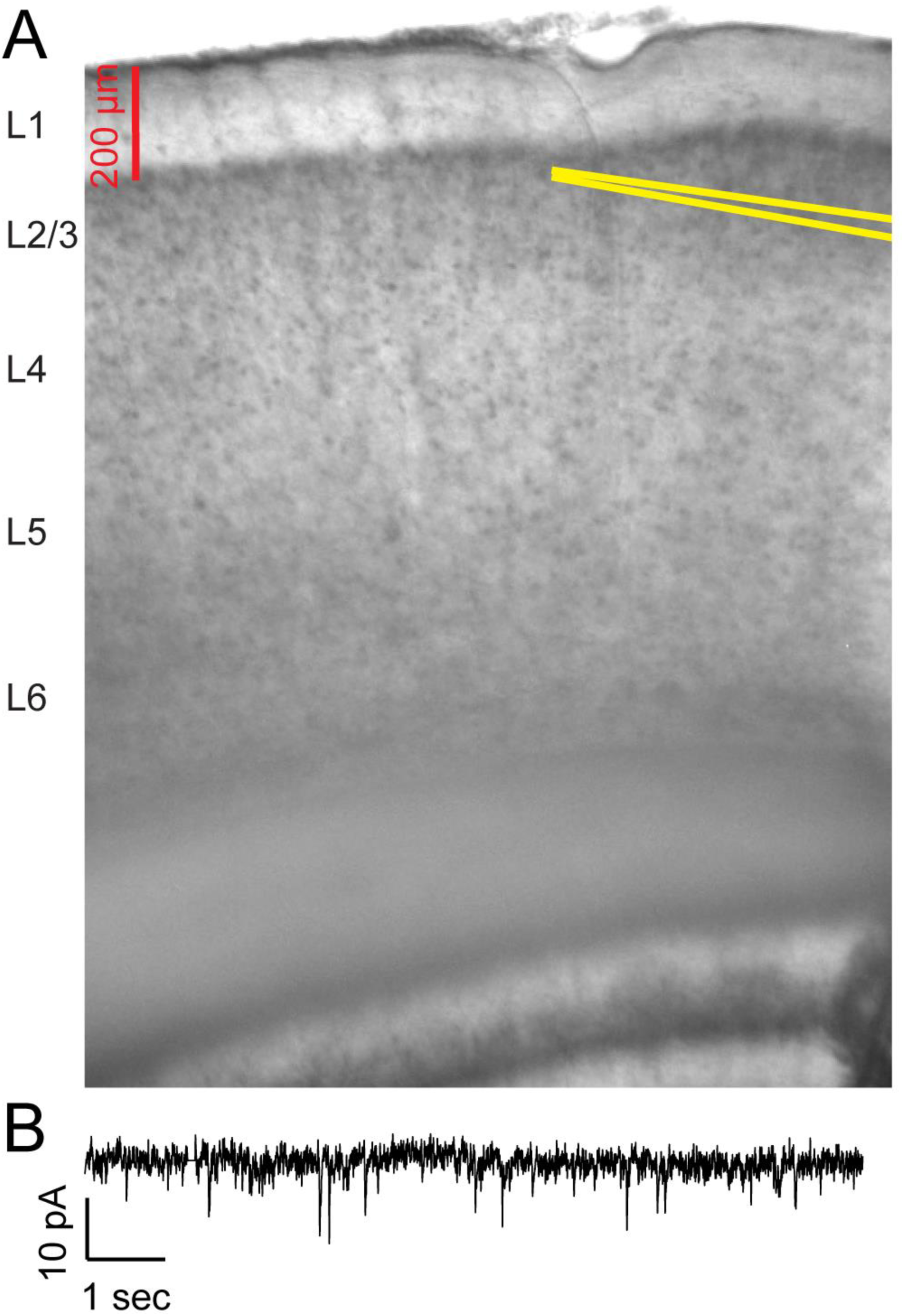
Image of auditory cortex slice and example voltage-clamp recording A. Example image of recording electrode placement in a coronal slice containing auditory cortex. Recording pipette walls are highlighted in yellow lines. B. Example of membrane current recorded in voltage clamp with holding potential of 10 mV and bath application of 20 μM GABAzine.

While holding the cell in voltage clamp, we recorded spontaneous excitatory and inhibitory postsynaptic currents (sEPSCs and sIPSCs), and miniature excitatory and inhibitory postsynaptic currents (mEPSCs and mIPSCs). IPSCs were recorded with bath application of 20 μM DNQX and 10 μM CPP, while the membrane potential was clamped at 10 mV with a Multiclamp 700B amplifier. EPSCs were recorded with bath application of 20 μM GABAzine while the membrane potential was clamped at −65 mV. Spontaneous currents were initially recorded for 15 minutes. Subsequently, 1 μM TTX was added to the aCSF, and miniature postsynaptic currents were recorded for the next 15 minutes. IPSC and EPSC events were detected and quantified using Minianalysis software. To determine the input resistance of the membrane, we periodically injected hyperpolarizing voltage pulses (−10 mV, 100 ms, 1 pulse per 20 seconds), and measured the median current response during the first five minutes of recording.

### Laser Scanning Photostimulation

MNI-caged-L-glutamate (Tocris) releases glutamate with exposure to 300-380 nm light, allowing for temporally and spatially precise uncaging of glutamate with laser photostimulation. We produced spatial maps of input strength in coronal slices of auditory cortex, prepared in the same manner as for spontaneous and miniature postsynaptic currents. To generate input maps, we applied to 150 μM MNI-glutamate to the aCSF. UV laser light (355 nm, 100 kHz pulses, DPSS Lasers) was guided to the slice by optical path mirrors and lenses (Thorlabs, Newport), and s focused through a 10x objective (Olympus). The beam intensity was attenuated with an acousto-optical modulator (Gooch and Housego), to deliver 24 mW light at the slice in 1 ms pulses. At this power, laser photostimulation was observed to drive spikes in neurons within a ∼50 μm radius around the laser spot center, similar to what has been seen using a similar laser stimulation configuration in the mouse auditory cortex (Slater et al. 2019)

Maps of synaptic input amplitude and charge to layer 2/3 neurons were produced by laser scanning photostimulation. 50 µM QX-314 was added to the internal solution to block voltage-gated sodium channels, and a neuron from layer 2/3 was patched in whole-cell configuration and recorded in voltage clamp. In a subset of experiments, two neurons from layer 2/3 were simultaneously patched and recorded during photostimulation. Using Prairie View software or ePhus, the slice was serially photostimulated in a 32×32 grid of stimulation sites, with adjacent grid points separated by a 40 μm distance, serving as a lower bound on the spatial resolution of our analysis. The grid was aligned to the pial surface of the slice, and oriented along the point on the surface closest to the patched neuron(s). Laser stimulation was pulsed for 1 ms at each stimulation site, and advanced to the successive stimulation site every second. The sequence of stimulation sites was arranged in a non-neighbor order. Patched neurons were recorded in voltage clamp held at −65 mV during the 32×32 photostimulation sequence to produce maps of excitatory input, then recorded in voltage clamp held at 10 mV during the photostimulation sequence to produce maps of inhibitory input.

### Laser Scanning Photostimulation Analysis

We measured the charge, amplitude, and latency of the current response to photostimulation at each site. Current signals recorded during photostimulation were lowpass filtered at 150 Hz. The baseline current, measured as the median during the 100 ms window preceding the laser onset, was subtracted from the signal, so that all measures are relative to the baseline current. All measures were computed from a 200 ms analysis window starting at the laser onset. Charge was computed by rectifying the current (positive rectification for inhibitory charge, and negative rectification for excitatory charge), and integrating the rectified current during 200 ms analysis window. Amplitude was measured as the maximum current for inhibitory input, and minimum current for excitatory current. Latency was measured as the first time point of the analysis window in which the current exceeded 10% amplitude. Current responses qualified as “significant” if their amplitudes exceeded 10 times the standard deviation of the baseline window current measured during the 100 ms prior to laser onset.

Maps of input charge and amplitude were constructed by ordering response charge and amplitude measures in two-dimensional arrays according to the photostimulation site. Observed amplitude and charges may include spontaneous currents in the recorded neuron that coincide with photostimulation. We took two approaches to reduce noise introduced by photostimulation-independent currents. First, we smoothed amplitude and charge maps by convolving the maps with a 4×4 gaussian kernel with standard deviation of 0.5. A similar approach was employed by Kratz and Manis (2015). Second, we included only charges and amplitudes from sites with significant responses. Thus, group-averaged maps and means were computed by including only significant sites from smoothed maps.

Excitatory responses to glutamate uncaging may arise from two sources. EPSCs may be driven by either: 1) the binding of photo-uncaged glutamate to ionotropic glutamate receptors in the recorded cell membrane, considered “direct” responses, or 2) synaptic transmission from presynaptic neurons driven by uncaged glutamate, considered “synaptic” responses. Synaptic responses were separated from direct responses on the basis that synaptic response latencies are later than 7 ms, and direct responses latencies are earlier than 7 ms. Response latencies were segregated at 7 ms because the distribution of synaptic latencies revealed a local minimum at 7 ms, indicating two subpopulation of EPSCs. We interpreted the subpopulation with earlier latencies as direct currents; therefore, we included in our group-averaged excitatory maps and means, only responses with latencies later than 7 ms, to capture synaptic responses. Comparable time windows of direct and synaptic responses have been observed by other studies (Kratz and Manis, 2015; Meng et al., 2017).

### Statistical Analysis

We tested the effect of PCB treatment on hearing thresholds, spontaneous synaptic current amplitude and frequency, and photostimulation-evoked synaptic current amplitude, charge, latency, input area, mean input distance, and excitation-inhibition ratios. However, these responses may vary with changes of age and hearing thresholds. Furthermore, we typically recorded from multiple neurons from each subject, and sampled multiple subjects from each litter (spontaneous and miniature currents: 116 neurons, 65 subjects, 24 litters, photostimulation-evoked currents: 46 neurons, 32 subjects, 26 litters), potentially introducing litter effects when comparing the responses of individual neurons. To account for response variance introduced by these factors, we used mixed effects modeling to predict our responses with PCB treatment, age, and noise hearing thresholds as fixed effects and birth litter as a grouping variable for random effects. For all response variables tested, we did not find a significant relationship between the response variables and age or hearing threshold. We report the significance of group comparisons for each response variable, as the probability of the slope of the response over PCB treatment, against a student’s t distribution.

Ratios of excitatory to inhibitory charge and amplitude are expressed as a gain in dB, as the logarithmic transform of the observed ratios approximately follows a gaussian distribution.

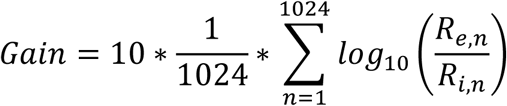

Here, R_e,n_ and R_i,n_ are the inhibitory and excitatory charge or amplitude response to photostimulation at site n (total stimulation sites = 1024), respectively.

To quantify differences in the spatial profiles of synaptic input, input charge was binned by stimulation site distance in 80 μm bins. Mean charge within each bin, across all subjects in each treatment group, was computed. Connection probability was computed as the number of sites with significant responses within each 80 μm bin, divided by the total number of sites within that bin.

## Results

### Hearing thresholds are elevated for PCB-exposed subjects

Two separate cohorts of rats were exposed to either a PCB mixture (6 mg/kg/day), or a corn oil vehicle, starting at gestation and continuing until weaning. We recorded ABRs to assess hearing thresholds during adulthood (110-518 days), and used a mixed-effects model to test the effect of PCB exposure on hearing threshold, independent of age and litter effects. A modest but significant hearing loss was seen with developmental PCB exposure, consistent with previous studies (Powers et al., 2006; Powers et al., 2009). We observed these hearing threshold differences in both rats used for comparing spontaneous and miniature synaptic currents (study 1), and those used to compare photostimulation-evoked currents (study 2). PCB-exposed rats had on average 9.0 dB higher ABR thresholds to white noise bursts relative to controls in the first study (t(33) = 6.86, p < 0.001), and 5.8 dB higher thresholds in the second (t(29) = 4.31, p < 0.001). ABRs to tone pips revealed that PCB exposure elevated thresholds to 4 kHz and 8 kHz tones in both studies, and elevated thresholds to 16 kHz tones in the second study (Fig. 2). Therefore, we confirmed that the PCB-exposed subjects in our study had elevated hearing thresholds, consistent with previous findings.

### Spontaneous and miniature inhibitory postsynaptic currents are more frequent with PCB exposure

We asked whether developmental PCB exposure would change inhibitory and excitatory synaptic input to the auditory cortex. To answer this question, we patched Layer 2/3 neurons from coronal slices of the auditory cortex and measured spontaneous excitatory or inhibitory postsynaptic currents in separate sets of recordings (see Fig. 3 for an example of slice image and spontaneous EPSCs). On average, cortical neurons in PCB-exposed subjects received more frequent (t(38) = 3.83, p < 0.001) and larger amplitude (t(38) = 2.70, p = 0.010) spontaneous IPSCs for PCB-exposed subjects compared to controls (Fig. 4A). In contrast, no difference was apparent in the frequency or amplitude of spontaneous EPSCs between the two treatment groups. To clarify the potential mechanisms of the synaptic changes, we isolated miniature synaptic currents with bath application of 1 μM TTX. Consistent with our observations of spontaneous synaptic currents, PCB treatment was associated with higher miniature IPSC frequency (t(28) = 2.66, p = 0.013), and not associated with differences in miniature EPSC frequency or amplitude. However, the amplitude of miniature IPSCS was not different between treatment groups (t(28) = 1.59, p = 0.12), suggesting that changes in inhibition with PCB exposure may be mediated primarily by presynaptic changes. Furthermore, input resistance was not affected by PCB exposure (PCB exposed: 188.2 ± 34.9 MΩ, control: 225.4 ± 41.7 MΩ, t(66) = 1.29, p = 0.21), suggesting that the increased spontaneous IPSC amplitudes are mediated by synaptic mechanisms, rather than changes in the intrinsic membrane properties of cortical neurons. Together, these data point to an increase of spontaneous inhibitory input to Layer 2/3 auditory cortex in PCB-exposed subjects.

**Figure 4.**
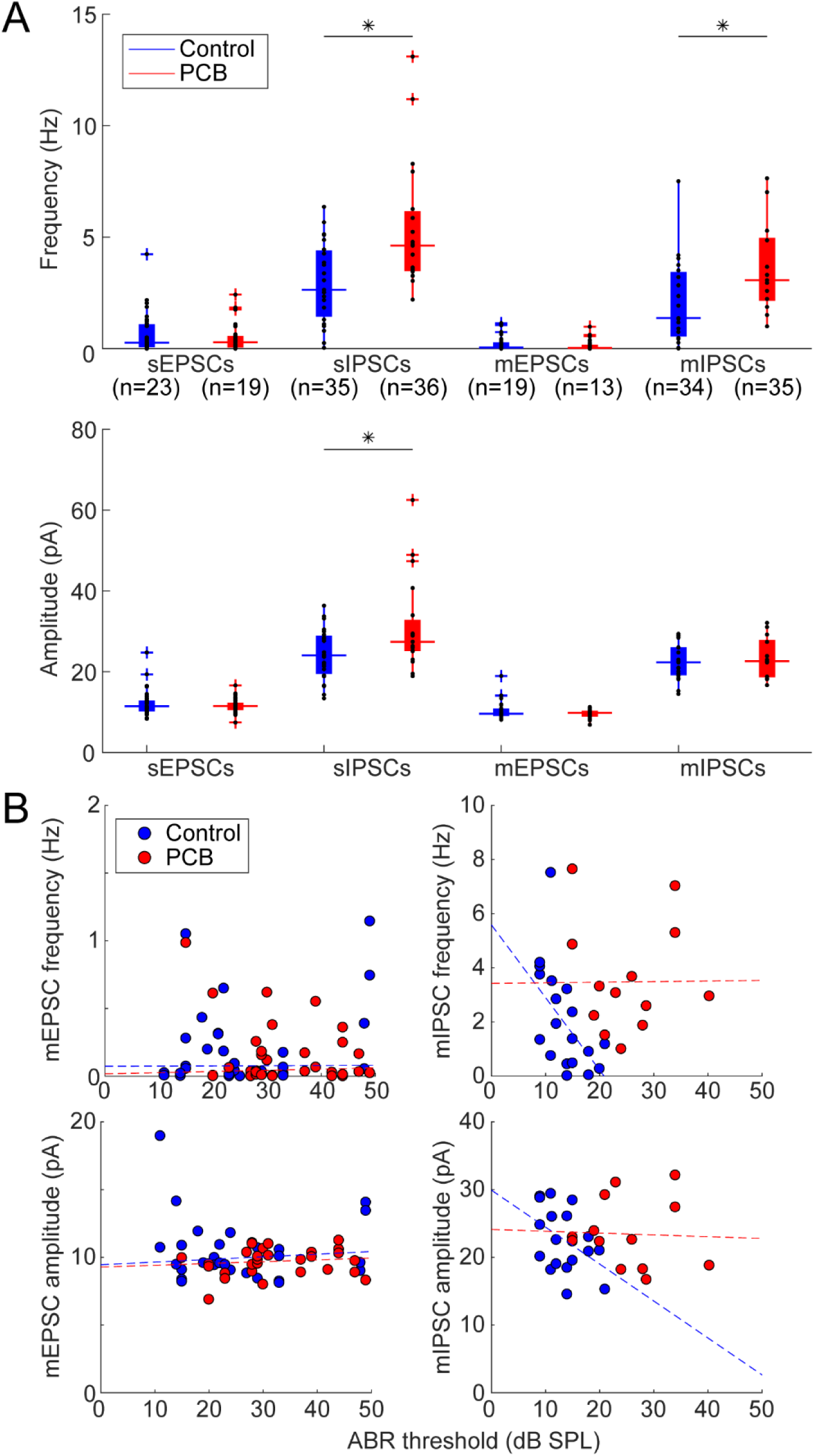
Comparison of spontaneous and miniature synaptic currents A. Comparison of frequency of synaptic currents between control (blue) and PCB (red) treatments. Boxplots indicate median (horizontal bar), 25^th^ and 75^th^ percentiles (box), range of non-outlier points (vertical whiskers), and outliers (crosses). Black asterisks indicate significant comparisons, * p < 0.05. B. Relationship of ABR threshold and sIPSC frequency for control (blue points) and PCB-exposed (red points) groups. Dashed lines indicate robust linear regression fits.

Changes of inhibitory and excitatory input to auditory cortex following peripheral hearing loss has been well documented (Kotak et al., 2005; Sarro et al., 2008; Balaram et al. 2019). Therefore, changes of inhibition seen in exposed subjects could be a secondary effect of the hearing threshold differences induced by PCB exposure. Among control subjects, increases of hearing threshold predicted reductions of sIPSC frequency (r = −0.44, p = 0.04) and amplitude (r = −0.42, p = 0.04, Fig. 4B, blue points), and reductions of mIPSC frequency (r = −0.58, p = 0.01). These data indicate that in control subjects, the auditory system can adjust cortical inhibition to the hearing sensitivity of the animal. However, this relationship was abolished in PCB-exposed subjects (sIPSC frequency: r = 0.15, p = 0.55, amplitude: r = −0.13, p = 0.59, mIPSC frequency: r = −0.02, p = 0.94, mIPSC amplitude: r = −0.03, p = 0.92), suggesting that the PCB-induced increases in cortical inhibition are not caused by peripheral hearing loss. In contrast to the relationship of hearing thresholds and synaptic inhibition, hearing thresholds did not correlate with sEPSC frequency in either control or PCB-exposed rats (control: r = 0.007, p = 0.97 PCB: r = 0.10, p = 0.55), amplitude (control: r = −0.03, p = 0.84, PCB: r = 0.22, p = 0.21), or mEPSC frequency (control: r = 0.29, p = 0.11 PCB: r = −0.21, p = 0.21) or amplitude (control: r = −0.052, p = 0.79 PCB: r = 0.17, p = 0.39).

### Laser scanning photostimulation reveals maps of synaptic input to Layer 2/3 auditory cortical neurons

Laser scanning photostimulation of caged glutamate (LSPS) allows spatially and temporally precise stimulation of the slice. By using LSPS during recordings of photostimulation-evoked synaptic currents, spatial maps of synaptic strength can be generated (Fig. 5B, C). To further elucidate changes in auditory cortical connectivity associated with PCB exposure, we examined the spatial profiles of synaptic input to layer 2/3 auditory cortex in exposed and control subjects. Fig. 6 summarizes the spatial maps produced with LSPS. Here we align all spatial maps to the recorded neuron, with the pial surface in the positive y direction, and dorsomedial in the positive × direction. Significant responses from stimulation site with the same positions relative to the recorded neuron are averaged across all neurons from each treatment group, and the averaged responses from each relative position are combined to produce the spatial maps. In excitatory maps, responses with a latency < 7ms are excluded, to minimize the influence of direct responses to glutamate.

**Figure 5.**
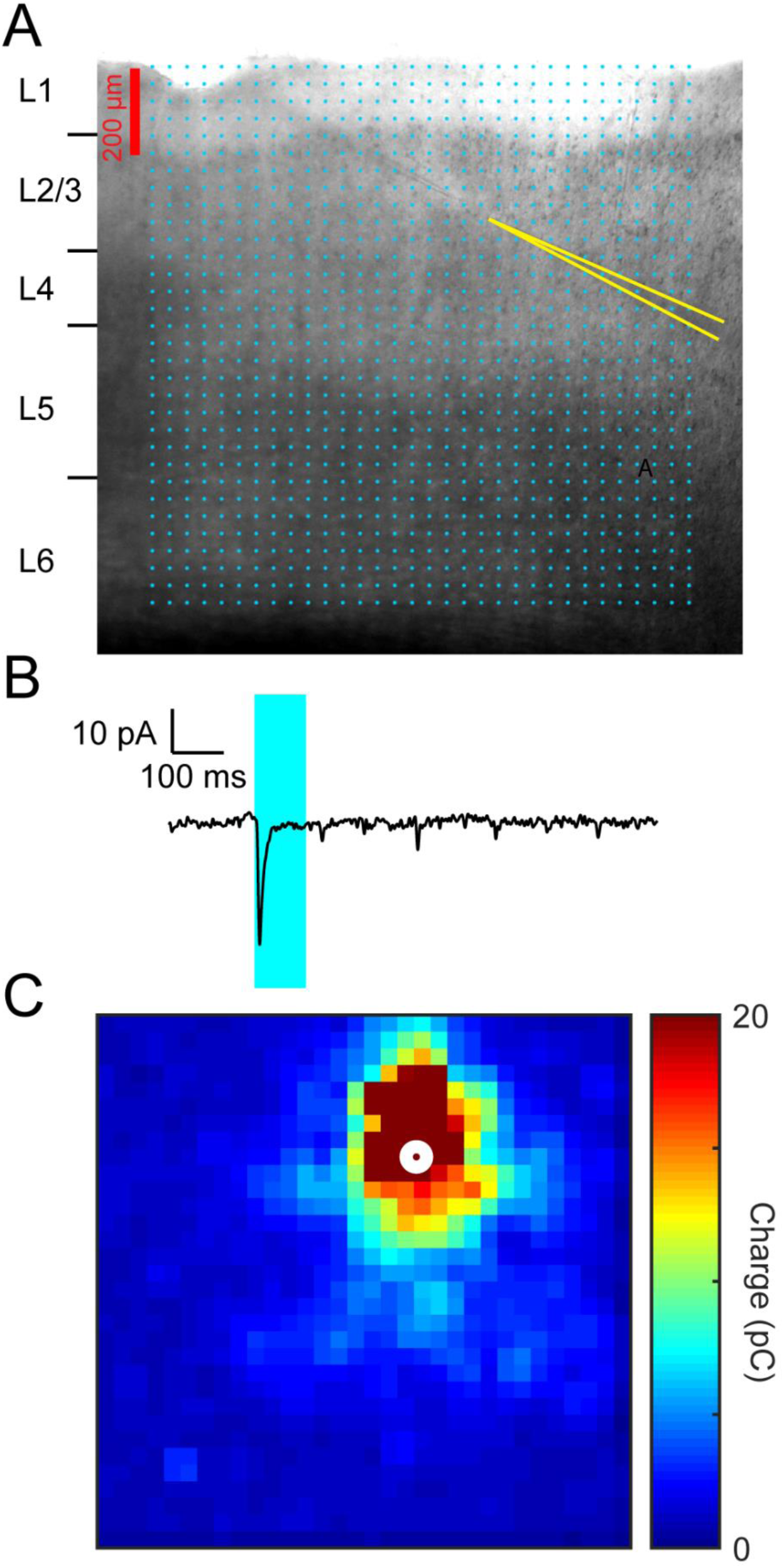
Demonstration of laser scanning photostimulation mapping of input charge A. Example image demonstrating positions of stimulation grid and recording electrodes in a coronal slice containing auditory cortex. Cyan points mark the sites of the 32 × 32 photostimulation grid. Recording pipette walls are highlighted in yellow lines. In the example, current recordings were simultaneously collected from two neurons. B. An example photostimulation-evoked current response from the cell positioned on the bottom right. Holding potential was −65 mV. Timing of the laser pulse (1 ms duration) is indicated by the red arrowhead. A pronounced negative peak begins shortly after the laser onset. C. Map of input charge from current responses to photostimulation at all sites of the 32 × 32 stimulation grid. For recordings of excitatory responses, measured charge is inverted to positive values, and represented by color.

**Figure 6.**
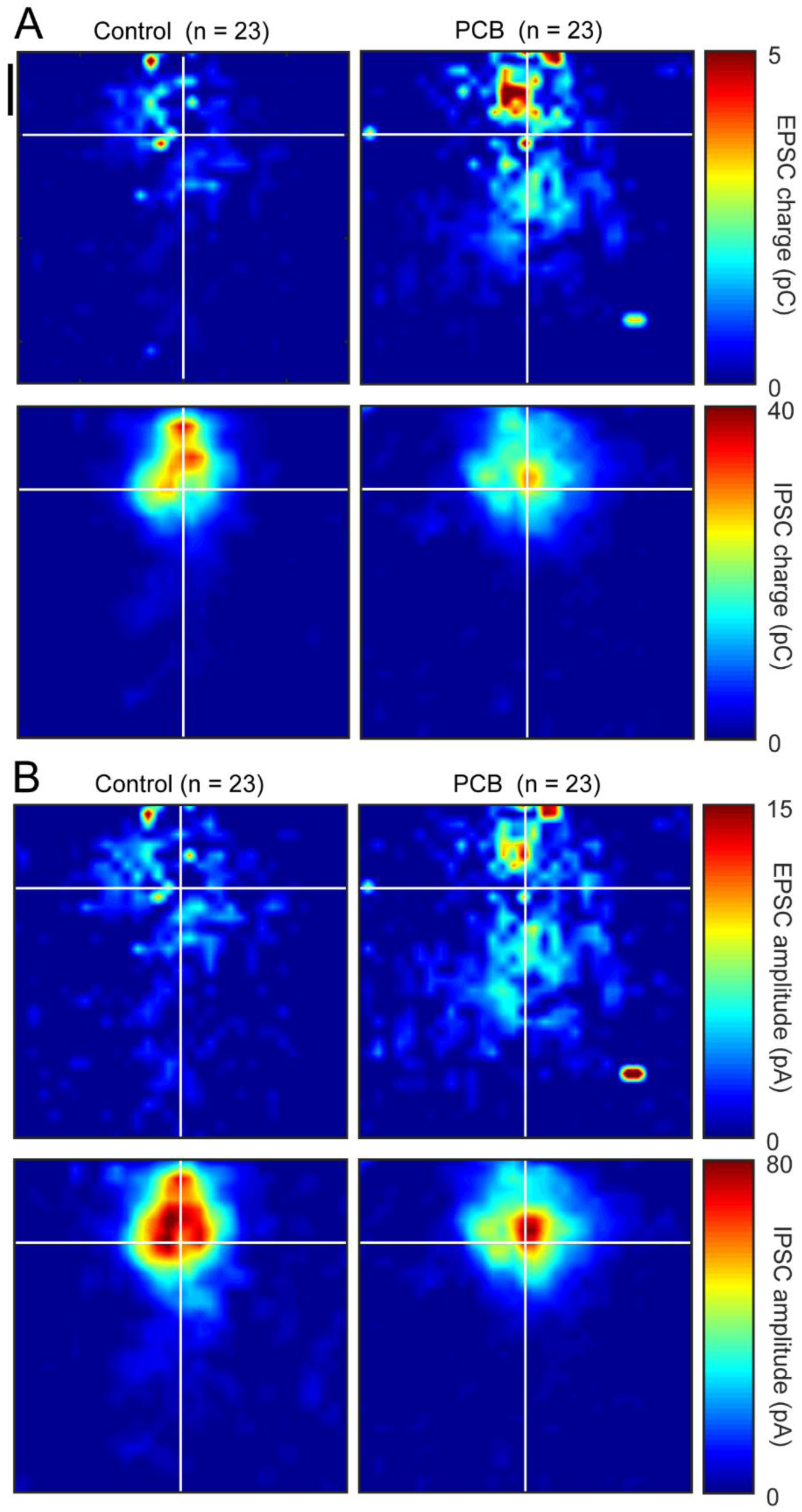
Group averaged maps of photostimulation-evoked synaptic strength Photostimulation-evoked input maps of charge (A), and amplitude (B) aligned to the recorded cell body. Measured charge at each site is averaged across all cells from each treatment group and is represented in color. EPSC charge maps are presented in the top plots, and IPSC charge maps are presented in the bottom plots. Black vertical scale bar marks 200 μm.

Among maps from control subjects, the strongest excitatory input (depicted in red pixels in Fig. 6) is on average near or superficial to the recorded neuron, with minimal excitatory input (depicted in dark blue pixels) from any point more than 400 μm distance from the recorded cell. In maps from PCB-exposed subjects, the strongest excitatory input is also near or superficial to the recorded neuron, with moderate input (light blue to cyan pixels) spanning most of the photostimulation map. Inhibitory maps for both control and PCB-exposed subjects were similar in shape. The strongest inhibitory input was near the recorded neuron, and average inhibitory strength drops off rapidly with increasing distance.

### PCB-exposed cells receive more total excitatory input charge, and more distant excitatory inputs

We estimated the total synaptic input for each recorded cell by summing the amplitudes and charges evoked from all stimulation sites that yielded a significant response (Fig. 7A-B, first two columns). Cells from control subjects received 0.12 ± 0.03 nC of total excitatory charge and 2.07 ± 0.72 nC of total inhibitory charge. Cells from PCB-exposed subjects received 0.31 ± 0.10 nC of total excitatory charge and 2.10 ± 1.10 nC of total inhibitory charge. PCB exposure was associated with a significant increase in excitatory charge (t(39) = 2.26, p = 0.030), but no change in inhibitory charge (t(39) = 0.029, p = 0.98). However, the total of photostimulation-evoked excitatory and inhibitory current amplitudes were not different between control (excitation: 0.59 ± 0.20 nA, inhibition: 6.40 ± 2.22 nA) and PCB-exposed groups (excitation: 1.09 ± 0.36 nA, inhibition: 5.09 ± 2.28 nA, comparison of excitation: t(39) = 1.52, p = 0.14, inhibition: t(39) = 0.28, p = 0.78). The ratio of excitation to inhibition was not significantly different between treatments (Charge E/I, control: −2.1 ± 1.8 dB, PCB-exposed: −3.6 ± 1.2 dB, t(30) = 0.56, p = 0.58; Amplitude E/I, control: −0.7 ± 1.7 dB, PCB-exposed: −2.2 ± 0.9 dB, t(30) = 0.52, p = 0.61). In addition, latency was not affected by PCB treatment (excitatory latencies, control: 17.2 ± 6.4 ms PCB: 17.2 ± 9.7 ms, t(37) = 0.31, p = 0.76, inhibitory latencies, control: 12.4 ± 7.2 ms, PCB: 12.3 ± 5.9 ms, t(38) = 0.62, p = 0.54). In summary, relative to controls, PCB-exposed cells responded to the excitatory inputs evoked by photostimulation with greater charge, but similar amplitude, and responded similarly to inhibitory inputs.

**Figure 7.**
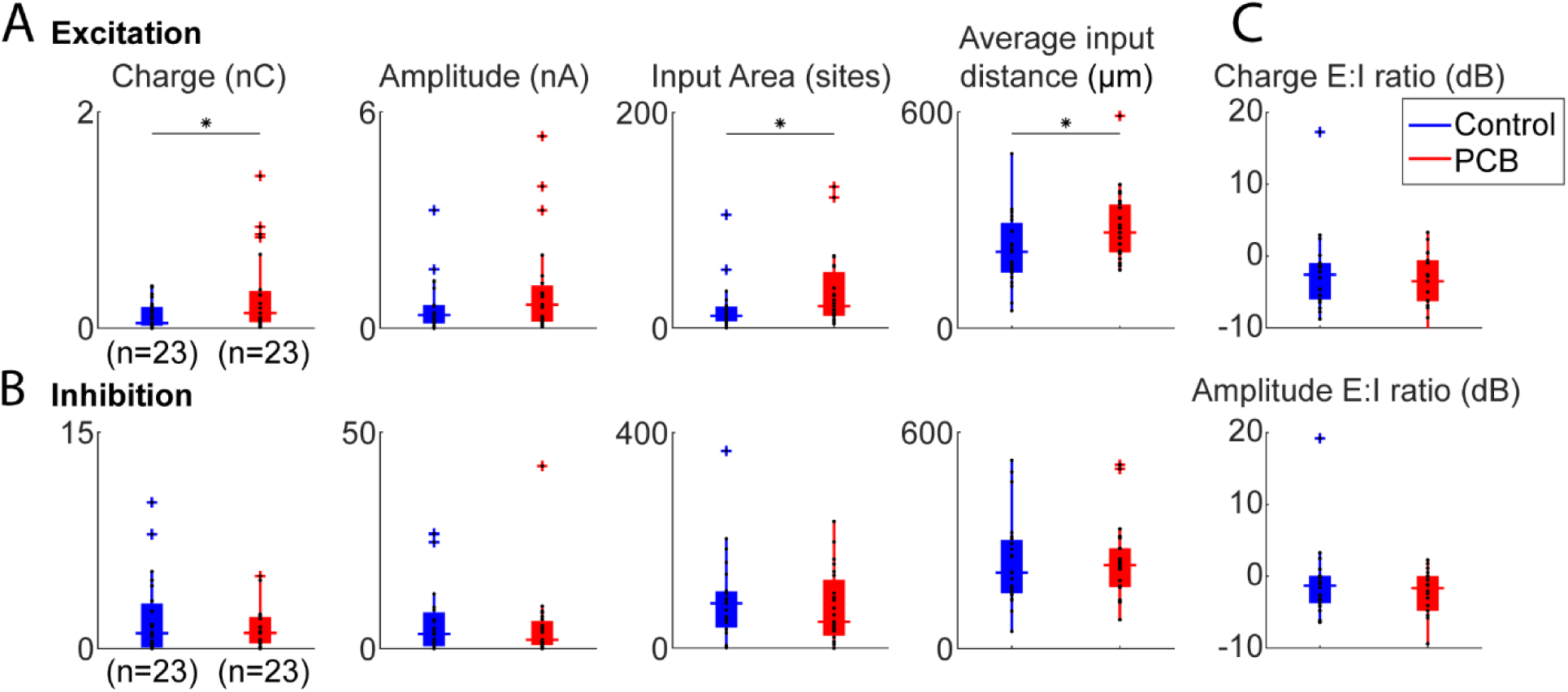
Comparison of photostimulation-evoked currents between treatments Comparison of total excitatory (A) and inhibitory (B) input charge, input area, and input distance, between control (blue) and PCB-exposed (red) treatment groups. Boxplots indicate median (horizontal bar), 25^th^ and 75^th^ percentiles (box), range of non-outlier points (vertical whiskers), and outliers (crosses). Black asterisks indicate significant comparisons, * p < 0.05. C. Ratio of excitatory to inhibitory charge between control and PCB-exposed groups.

The difference in total excitatory charge between PCB-exposed and control subjects may be due to a change in the number of input sites, or the strength of each input site. We observed that cells from PCB-exposed subjects received significant excitatory input from more sites than cells from controls (control: 17.9 ± 6.0 sites, PCB-exposed: 33.8 ± 9.3 sites, p = 0.048). On the other hand, the average charge per site was not significantly different between treatments (control: 7.2 ± 1.3 pA, PCB-exposed: 9.0 ± 1.9 pA, t(39) = 0.35, p = 0.72). Thus, cortical neurons from PCB-exposed rats receive input from a greater number of sites than controls but receive input of the same strength from each site.

The spatial profile of input strength is plotted as a function of stimulation site distance in Fig. 8. Differences in excitatory input response charge, and excitatory input site number are most pronounced between ∼100 and 700 μm distance. Furthermore, the average distance of significant excitatory stimulation sites is longer in the PCB-exposed group compared to controls (control: 215 ± 25.8 μm, PCB-exposed: 284 ± 25.3 μm, t(39) = 2.23, p = 0.032). In contrast, the average distance of significant inhibitory stimulation sites is not different between the groups (control: 239 ± 32 μm, PCB: 239 ± 29 μm, t(38) = 0.16, p = 0.87). Distances were clustered into layers based on contrast seen on DIC images, and no differences were seen between groups (not shown). Together, these findings suggest that in PCB-exposed subjects, Layer 2/3 auditory cortical neurons integrate excitatory input from a greater number of neurons at intermediate (100-700 µm) distances, without affecting inhibitory connections.

**Figure 8.**
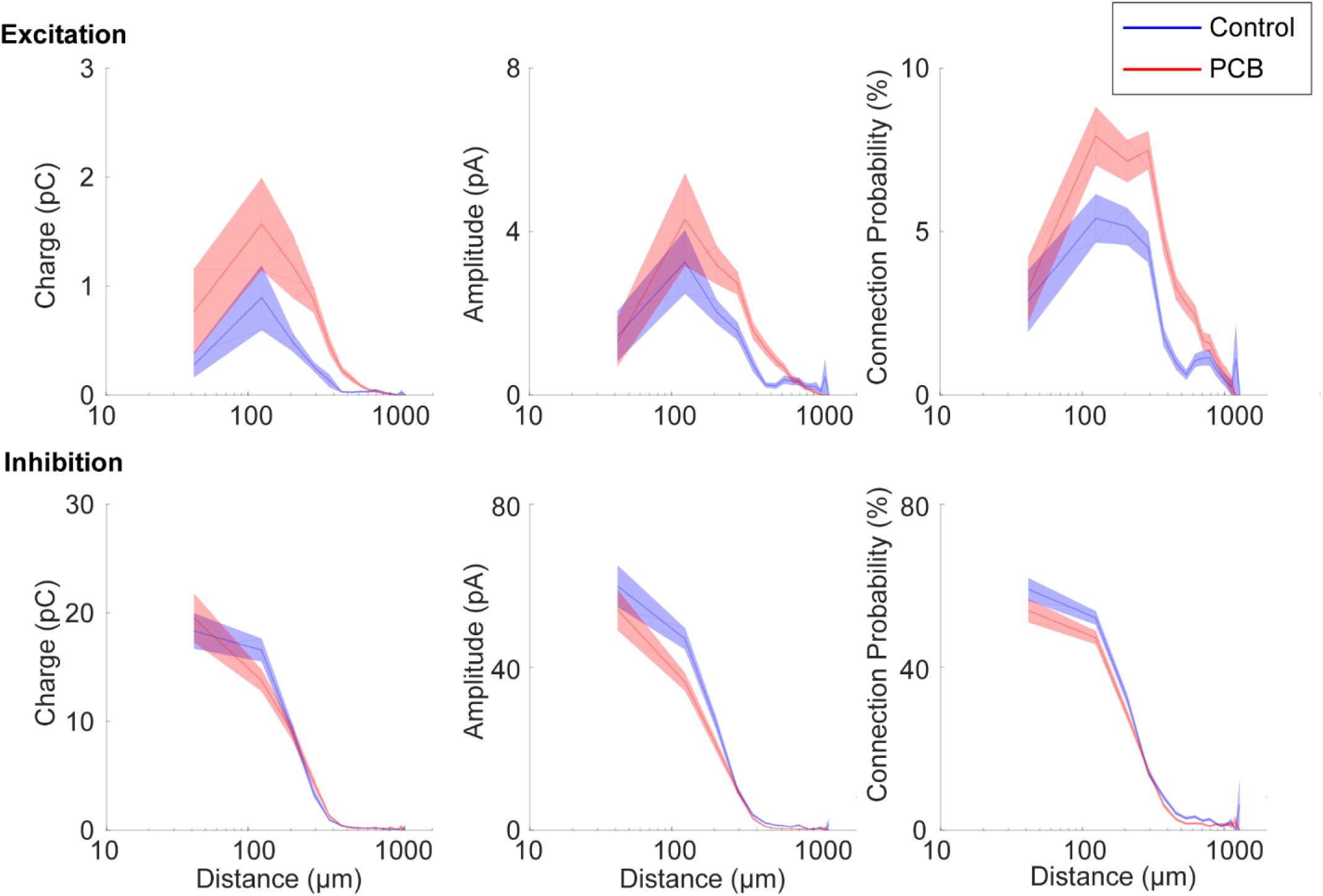
Spatial profile of synaptic input Distance profile of input charge and connection probability for control (blue) and PCB-exposed (red) treatment groups. Response measures are binned by input distance in 80 μm bins, and interpolation between bin means are marked by the solid lines. The shaded areas indicate 1 standard error bounds around the means.

## Discussion

### Developmental PCB exposure induces long-lasting increases in spontaneous inhibitory tone and increases in excitatory connectivity in auditory cortex

We found that developmental PCB exposure results in paradoxically increased spontaneous IPSC amplitude and frequency and increased miniature IPSC frequency in layer 2/3 of the auditory cortex (Fig. 4A), while also inducing peripheral hearing loss. The increased inhibition appears to be mediated by presynaptic changes, as the amplitude of miniature IPSCs is unchanged with PCB treatment. In contrast, PCB treatment did not affect photostimulation-evoked IPSCs. Therefore, PCB exposure may induce changes that specifically affect spontaneous release of GABA from inhibitory cortical neurons, without affecting the strength of evoked inhibitory synaptic transmission. While evoked synaptic currents depend on synaptic density and efficacy of each synapse, spontaneous synaptic transmission reflects synaptic efficacy and rate of vesicle release. Therefore, PCB exposure may increase vesicle release of inhibitory inputs. Vesicle release can be modified by a variety of changes, including changes of activity, membrane potential, intracellular calcium dynamics, and the readily releasable pool. Increases of spontaneous inhibition following developmental PCB exposure are consistent with previous findings that thalamocortical transmission is more strongly enhanced by GABA_A_-receptor blockade in PCB-exposed subjects (Sadowski et al., 2016). Because PCB exposure increases spontaneous inhibitory input to auditory cortical neurons, release from inhibition with GABA_A_-receptor antagonist application is more pronounced in PCB-exposed subjects compared to controls.

It is important to note that rats used to study photostimulation-evoked currents had unexpectedly higher white noise thresholds and lower pure tone thresholds than those used to study spontaneous and miniature currents (t(106) = 6.84, p < 0.001). The reasons for these differences are not known, but may include: 1) evaluation of ABR thresholds by different experimenters, 2) different headstage and preamplifiers used between studies (study 1: Dagan 2400A, study 2: TDT RA4LI/RA4PA), or 3) true hearing threshold differences between the different cohorts of rats, purchased approximately two years apart. Nonetheless, PCB exposed rats showed significantly higher noise, 4 kHz tone, and 8 kHz tone thresholds in each study.

Developmental PCB exposure does not affect the frequency and amplitude of spontaneous or miniature EPSCs (Fig. 4A), suggesting that synaptic transmission of individual synapses and spontaneous excitatory input do not change with PCB exposure. However, in neurons from PCB-exposed subjects, EPSCs were evoked from a larger number of photostimulation sites, and they were evoked from more distant sites on average (Fig. 7A). Thus, rather than changing the strength of individual synapses, PCB exposure results in abnormally enhanced connectivity between excitatory cortical neurons, with the largest changes at distances of 100-700 μm (Fig. 8). Because Layer 2/3 cortical neurons integrate input from neurons with different frequency tuning, PCB-induced changes in excitatory connectivity may disrupt frequency receptive fields in auditory cortex (Kenet et al., 2006). Together, these changes suggest that PCB exposure may degrade spectral resolution as a result of excessive excitatory connectivity in the cortex.

Among the control subjects, higher ABR thresholds are associated with reduced cortical inhibition, which may reflect a compensatory increase of gain in the central auditory system (Fig. 4B). However, with PCB exposure, inhibition is not related to hearing threshold. Therefore, PCB exposure may disrupt the compensatory regulation of cortical activity by modulation of central auditory gain seen in unexposed subjects. Furthermore, PCB exposure increases spontaneous inhibitory input in the cortex, while elevating hearing thresholds. Thus, while PCB exposure impairs hearing, its effects on spontaneous inhibition would paradoxically further reduce activity in the auditory cortex, potentially compounding auditory perceptual deficits.

### Potential mechanisms of changes

Increases of spontaneous IPSC amplitude and frequency seen in PCB-exposed subjects were unexpected and contrast previous findings of reduced central inhibition following hearing loss (Bledsoe et al., 1995; Vale and Sanes, 2002; Kotak et al., 2005; Sarro et al., 2008; Balaram et al., 2019). Several differences between PCB-induced hearing loss and aforementioned studies may explain differences in the outcomes. First, the hearing impairment following PCB exposure is relatively mild, elevating thresholds by less than 10 dB on average as observed in the current study and in previous studies (Powers et al., 2006; Powers et al., 2009). Exposure to PCBs leads to a loss of outer hair cells and reduced otoacoustic emissions (Goldey et al., 2000; Lasky et al., 2002; Powers et al., 2006; Trnovec et al., 2008). Inner hair cells, on the other hand, are spared after exposure to a commercial PCB mixture (Aroclor 1254, Goldey et al., 2000), but it is not yet known if they are affected by the Fox River PCB mixture. PCBs and other dioxin-like compounds reduce thyroid hormone levels, and this thyroid hormone deficiency may be involved in PCB-induced hearing loss, as thyroxine replacement partially restores hearing in PCB-exposed animals (Goldey et al., 1995, 1998; Poon et al., 2011). Furthermore, developmental hypothyroidism can affect the development, connectivity, and organization of auditory cortical neurons (Ruiz-Marcos et al., 1983; Berbel et al., 1993; Lucio et al., 1997). In contrast, studies documenting increases of central auditory gain following hearing loss typically involve damage to inner hair cells and threshold increases of 30 dB or more. Therefore, PCB-induced hearing loss may involve specific mechanisms not typically seen with peripheral hearing loss, that result in increased cortical inhibition.

Second, in addition to damaging the sensory epithelium, PCBs also have direct actions in the central nervous system. The Fox River PCB mixture used in this study was previously found to increase binding of ryanodine to ryanodine receptors (RyRs, Kostyniak et al., 2005). Ryanodine receptor activation can induce growth of dendrites, and RyR-dependent increases in dendritic growth have been observed following developmental exposure to PCBs (Lein et al., 2007; Yang et al, 2009). Furthermore, developmental PCB exposure disrupts both experience-dependent synaptic plasticity and Morris water maze learning, supporting the idea that the activation of RyRs may underlie some of the behavioral effects of PCBs (Yang et al., 2009). PCB exposure not only affects dendritic growth, but also alters excitatory and inhibitory synaptic transmission in auditory cortex and hippocampus (Kenet et al., 2007; Kim et al., 2009). In hippocampal slices, changes in synaptic transmission following wash-in of PCBs were found to be dependent on RyR activation (Kim et al., 2009). We observed an increase in the number of sites producing significant excitatory synaptic responses to laser photostimulation (Fig. 7A, 8). This change may be explained by increased synaptic connectivity between excitatory cortical neurons due to higher levels of RyR activation in PCB-exposed subjects.

In summary, we find that developmental exposure to PCBs increases spontaneous inhibitory input to the neurons in layer 2/3 of the auditory cortex, increases the number of excitatory connections, and disrupts the relationship between inhibition and hearing impairment. These changes were unexpected as the auditory system typically responds to hearing loss by increasing gain in central auditory structures. Thus, in addition to elevating hearing thresholds, PCB exposure may disrupt plastic changes needed to restore central auditory function after hearing loss by increasing spontaneous cortical inhibition. Thus, the cognitive deficits associated with PCB exposure in humans (Vreugdenhil et al., 2002; Schantz et al., 2003; Newman et al., 2006), may be related to long-lasting changes in the underlying synaptic architecture that alter local cortical network connectivity that are due to direct effects of PCBs on the brain.

## Acknowledgements

This work was supported by the National Institute of Environmental Health Sciences (NIEHS-R01 ES015687 to S.L.S., NIEHS-T32 ES007326 to C.M.L.), a Beckman Institute Postdoctoral Fellowship (R.N.S.) and funding from the University of Illinois Research Board. We thank Alex Asilador for assistance with speaker calibration, and Mindy Howe for help with dosing and breeding.

## Notes

Conflict of Interests: The authors declare no competing financial interests.

### Competing Interest Statement

The authors have declared no competing interest.

## References

Agency for Toxic Substances and Disease Registry (2000) Toxicological profile for polychlorinated biphenyls (PCBs). US Dept Health Services, Public Health Service.

Atencio CA, Schreiner CE (2010) Columnar connectivity and laminar processing in cat primary auditory cortex. PLOS ONE 5:1–18

Balaram P, Hackett TA, Polley DB (2019) Synergistic transcriptional changes in AMPA and GABA_A_ receptor genes support compensatory plasticity following unilateral hearing loss. Neurosci 407:108–119

Bandara SB, Eubig PA, Sadowski RN, Schantz SL (2016) Developmental PCB exposure increases audiogenic seizures and decreases glutamic acid decarboxylase in the inferior colliculus. Toxicol Sci 149:335–345

Berbel P, Guadaño-Ferraz A, Martínez M, Quiles JA, Balboa R, Innocenti GM (1993) Organization of auditory callosal connections in hypothyroid rats. Eur J Neurosci 5:1465–1478.

Bledsoe SC, Nagase S, Miller JM, Altschuler RA (1995) Deafness-induced plasticity in the mature central auditory system. Neuroreport 7:225–229

Chambers AR, Resnik J, Yuan Y, Whitton JP, Edge AS, Liberman MC, Polley DB (2016) Central gain restores auditory processing following near-complete cochlear denervation. Neuron 89:867–879

Crinnion WJ (2011) Polychlorinated biphenyls: persistent pollutants with immunological, neurological, and endocrinological consequences. Altern Med Rev 16:5–13

Crofton KM, Ding DL, Padich R, Taylor M, Henderson D (2000) Hearing loss following exposure during development to polychlorinated biphenyls: A cochlear site of action. Hear Res 144:196–204.

Goldey ES, Crofton KM (1998) Thyroxine replacement attenuates hypothyroxinemia, hearing loss, and motor deficits following developmental exposure to Aroclor 1254 in rats. Toxicol Sci 45:94–105.

Goldey ES, Kehn LS, Lau C, Rehnberg GL, Crofton KM (1995) Developmental exposure to polychlorinated biphenyls (Aroclor 1254) reduces circulating thyroid hormone concentrations and causes hearing deficits in rats. Toxicol Appl Pharmacol 135:77–88

Grandjean P, Weihe P, Burse VW, Needham LL, Storr-Hansen E, Heinzow B, (2001) Neurobehavioral deficits associated with PCB in 7-year-old children prenatally exposed to seafood neurotoxicants. Neurotoxicol Teratol 23(4):305–317

Jiang X, Wang G, Lee AJ, Stornetta RL, Zhu JJ (2013) The organization of two new cortical interneuronal circuits. Nat Neurosci 16:210–218

Kato HK, Gillet SN, Isaacson JS (2015) Flexible sensory representations in auditory cortex driven by behavioral relevance. Neuron 88:1027–1039

Kenet T, Froemke RC, Schreiner CE, Pessah IN, Merzenich MM (2007) Perinatal exposure to a noncoplanar polychlorinated biphenyl alters tonotopy, receptive fields, and plasticity in rat primary auditory cortex. Proc Nat Acad Sci 104:7646–7651.

Kim KH, Inan SY, Berman RF, Pessah IN (2009) Excitatory and inhibitory synaptic transmission is differentially influenced by two ortho-substittued polychlorinated biphenyls in the hippocampal slice preparation. Toxicol and App Pharmacol 237:168–177

Kostyniak PJ, Hansen LG, Widholm JJ, Fitzpatrick RD, Olson JR, Jelferich JL, Kim KH, Sabel HJK, Seegal RF, Pessah IN, Schantz SL (2005) Formulation and characterization of an experimental PCB mixture designed to mimic human exposure from contaminated fish. Toxicol Sci 88:400–411

Kotak VC, Fujisawa S, Lee FA, Karthikeyan O, Aoki C, Sanes DH (2005) Hearing loss raises excitability in the auditory cortex. J Neurosci 25:3908–3918

Kratz MB, Manis PB (2015) Spatial organization of excitatory synaptic inputs to layer 4 neurons in mouse primary auditory cortex. Front in Neural Circuits 9:1–17

Lasky RE, Widholm JJ, Crofton KM, Schantz SL (2002) Perinatal exposure to Aroclor 1254 impairs distortion product otoacoustic emissions (DPOAEs) in rats. Toxicol Sci 68:458–464

Lein PJ, Yang D, Bachstetter AD, Tilson HA, Harry GJ, Mervis RF, Kodavanti PRS (2007) Ontogenetic alternations in molecular and structural correlates of dendritic growth after developmental exposure to polychlorinated biphenyls. Env Health Perspect 115:556–563

Li MC, Wu HP, Yang CY, Chen PC, Lamber GH, Guo YL (2015) Gestational exposure to polychlorinated biphenyls and dibenzofurans induced asymmetric hearing loss: Yucheng children study. Environ Res 137:65–71

Lomber S, Malhotra SG (2007) Sound localization during homotopic and heterotopic bilateral cooling deactivation of primary and nonprimary auditory cortical areas in the cat. J Neurophys 97:26–43

Lucio RA, García-Velasco JV, Cerezo JR, Pacheco P, Innocenti GM, Berbel P (1997) The development of auditory callosal connections in normal and hypothyroid rats. Cereb Cortex 7:303–306

Meng X, Kao JPY, Lee HK, Kanold PO (2017) Intracortical circuits in thalamorecipient layers of auditory cortex refine after visual deprivation. eNeuro 4:1–11

Min JY, Kim R, Min KB (2014) Serum polychlorinated biphenyls concentrations and hearing impairment in adults. Chemosphere 102:6–11.

Newman J, Aucompaugh A, Schell LM, Denham M, DeCaprio AP, Gallo MV, Ravenscroft J, Kao CC, Hanover MR, David D, Jacobs AM, Tarbell AM, Worswick P, Akwesasne Task Force on the Environment (2006) PCBs and cognitive functioning of Mohawk adolescents. Neurotoxicol Teratol 28:439–45.

Noreña AJ (2011) An integrative model of tinnitus based on a central gain controlling neural sensitivity. Neurosci and Biobehav Rev 35:1089–1109

Oviedo HV, Bureau I, Svoboda K, Zador AM (2010) The functional asymmetry of auditory cortex is reflected in the organization of local cortical circuits. Nat Neurosci 13:1413–1420

Paxinos G, Franklin KBJ (2004) The mouse brain in stereotaxic coordinates. San Diego: Gulf Professional

Poon E, Bandara SB, Allen JB, Sadowski RN, Schantz SL (2015) Developmental PCB exposure increases susceptibility to audiogenic seizures in adulthood. NeuroTox 46:117–124.

Poon E, Powers BE, McAlonan RM, Ferguson DC, Schantz SL (2011) Effects of developmental exposure to polychlorinated biphenyls and/or polybrominated diphenyl ethers on cochlear function. Toxicol Sci 124:161–168

Powers BE, Widholm JJ, Lasky RE, Schantz SL (2006) Auditory deficits in rats exposed to an environmental PCB mixture during development. Toxicol Sci 89:415–422

Ruiz Marcos A, Salas J, Sanchez-Toscano F, Escobar del Rey F, Morreale de Escobar G (1983) Effect of neonatal and adult-onset hypothyroidism on pyramidal cells of the rat auditory cortex. Dev Brain Res 9:205–213.

Sadowski RN, Stebbings KA, Slater BJ, Bandara SB, Llano DA, Schantz SL (2016) Developmental exposure to PCBs alters the activation of the auditory cortex in response to GABA_A_ antagonism. NeuroTox 56:86–93

Sarro EC, Kotak VC, Sanes DH, Aoki C (2008) Hearing loss alters the subcellular distribution of presynaptic GAD and postsynaptic GABA_A_ receptors in the auditory cortex. Cereb Cortex 18:2855–2867

Schantz SL, Widholm JJ, Rice DC (2003) Effects of PCB exposure on neuropsychological function in children. Environ Health Perspect 111:357–376.

Slater BJ, Sons SK, Yudintsev G, Lee CM, Llano DA (2019) Thalamocortical and intracortical inputs differentiate layer-specific mouse auditory corticollicular neurons. J Neurosci 39:256–270

Sun W, Lu J, Stolzberg D, Gray L, Deng A, Lobarinas E, Salvi RJ (2009) Salicylate increases the gain of the central auditory system. Neurosci 159:325–334

Threlkeld SW, Penley SC, Rosen GD, Fitch RH (2008) Detection of silent gaps in white noise following cortical deactivation in rats. Neuroreport 19:893–898

Trnovec T, Šovčíková E, Hust’ák M, Wimmerová S, Kočan A, Jurečková D, Langer P, Palkovičová LU, Drobná B (2008) Exposure to polychlorinated biphenyls and hearing impairment in children. Envir Toxicol and Pharmacol 25:183–187

Vale C, Sanes DH (2002) The effect of bilateral deafness on excitatory and inhibitory synaptic strength in the inferior colliculus. Europ J Neuro 16:2394–2904

Vreugdenhil HJI, Lanting CI, Mulder PGH, Boersma ER, Weisglas-Kuperus N (2002) Effects of prenatal PCB and dioxin background exposure on cognitive and motor abilities in Dutch children at school age. J Pediatr 140:48–56.

Vreugdenhil HJI, Van Zanten GA, Brocaar MP, Mulder PGH (2004) Prenatal exposure to polychlorinated biphenyls and breastfeeding: opposing effects on auditory P300 latencies in 9-year-old Dutch children. Devel Med and Child Neurol 46:398–405

Wang J, Ding D, Salvi RJ (2002) Functional reorganization in chinchilla inferior colliculus associated with chronic and acute cochlear damage. Hear Res 168:238–249

Winkowski DE, Kanold PO (2013) Laminar transformation of frequency organization in auditory cortex. J Neurosci 33:1498–1508

Yang D, Kim KH, Phimister A, Bachstetter AD, Ward TR, Stackman RW, Mervis RF, Wisniewski AB, Klein SL, Kodavanti PRS, Anderson KA, Wayman G, Pessah IN, Lein PJ (2009) Developmental exposure to polychlorinated biphenyls interferes with experience-dependent dendritic plasticity and ryanodine receptor expression in weanling rats. Env Health Persp 117:426–435

Yang S, Su W, Bao S (2012) Long-term, but not transient, threshold shifts alter the morphology and increase the excitability of cortical pyramidal neurons. J Neurophysiol 108:1567–1574

Zeng F (2013) An active loudness model suggesting tinnitus as increased central noise and hyperacusis as increased nonlinear gain. Hear Res 295:172–179

Znamenskiy P, Zador TM (2013) Corticostriatal neurons in auditory cortex drive decisions during auditory discrimination. Nature 497:482–487

